# 20-HETE-promoted cerebral blow flow autoregulation is associated with enhanced α-smooth muscle actin positive cerebrovascular pericyte contractility

**DOI:** 10.1101/2021.01.20.427495

**Authors:** Yedan Liu, Huawei Zhang, Tina Yu, Xing Fang, Jane J. Ryu, Baoying Zheng, Zongbo Chen, Richard J. Roman, Fan Fan

## Abstract

We previously reported that deficiency in 20-HETE or CYP4A impaired the myogenic response and autoregulation of cerebral blood flow (CBF) in rats. The present study demonstrated that CYP4A was coexpressed with alpha-smooth muscle actin (α-SMA) in vascular smooth muscle cells (VSMCs) and most pericytes along parenchymal arteries (PAs) isolated from SD rats. Cell contractile capabilities of cerebral VSMCs and pericytes were reduced with a 20-HETE synthesis inhibitor, N-Hydroxy-N′-(4-butyl-2-methylphenyl)-formamidine (HET0016) but restored with 20-HETE analog 20-hydroxyeicosa-5(Z),14(Z)-dienoic acid (WIT003). Similarly, intact myogenic responses of the middle cerebral artery and PA of SD rats decreased with HET0016 and rescued by WIT003. Lastly, HET0016 impaired well autoregulated CBF in the surface and deep cortex of SD rats. These results demonstrate that 20-HETE has a direct effect on cerebral mural cell contractility that may play an essential role in CBF autoregulation.

## 1. Introduction

20-Hydroxyeicosatetraenoic acid (20-HETE) is an arachidonic acid metabolite by enzymes of CYP4A and CYP4F families. It has diverse physiological and pathological effects on cardiovascular function, implicating it may play a role in the pathogenesis, progression, and prognosis of various cardiovascular diseases [1-9]. 20-HETE contributes to sodium transport regulation, endothelial dysfunction, vasoconstriction, inflammation, angiogenesis, vascular remodeling, and restenosis. Alteration of 20-HETE production has been linked to hypertension; stroke; Alzheimer’s disease (AD); ischemia-reperfusion injury in the kidney, brain, and heart; as well as changes in airway and placenta vascular resistance [1, 10-14].

We have been studying the roles of 20-HETE in cerebral vascular function for decades. Our previous studies demonstrated that Dahl salt-sensitive (Dahl S) rats have a genetic deficiency in the production of 20-HETE and the expression of CYP4A enzymes [15]. They exhibited impaired myogenic response of the middle cerebral artery (MCA) and autoregulation of cerebral blood flow (CBF) [16]. This impaired cerebral vascular hemodynamics found in Dahl S rats is associated with enhanced blood-brain barrier (BBB) leakage in response to acute hypertension, which was rescued by knocking in wildtype *Cyp4a1* onto the Dahl S transgenic background [16]. On the other hand, previous studies demonstrated that inhibition of 20-HETE attenuated inflammation and oxidative stress in spontaneously hypertensive rats (SHR) in association with improvement of cerebral vascular function [9], and the enhanced cerebral vascular production of 20-HETE in this model was in association with ischemic stroke [17]. These results in terms of the role of 20-HETE and cerebral vascular function are controversial, possibly because different models with different genetic, anatomic, endocrine, and metabolic physiological or pathological conditions only display a “summarized” cerebral vascular phenotype. However, whether 20-HETE plays a direct role in cerebral vascular reactivity and the direction of its effect on the alternation of susceptibility to stroke and cognitive deficits have not been studied. In this regard, human studies demonstrated that genetic variants of *CYP4A11* and *CYP4F2*, the most potent enzymes producing 20-HETE in humans, have been linked to hypertension and stroke [18-29]. Several of these variants were validated *in vitro* that reduce the production of 20-HETE [24, 27, 30]. Furthermore, we previously reported that genetic variants in these two genes are associated with loss of cortical, hippocampal, and AD signature volumes and cognitive dysfunction in 4,286 elderly subjects in the Atherosclerosis Risk in Communities Neurocognitive Study (ARIC-NCS) [10].

20-HETE has been reported to be produced in cerebral vascular smooth muscle cells (VSMCs), endothelial cells (ECs), pericytes, astrocytes, neurons, and podocytes [31-36]. It has been well established that cerebral VSMCs play an essential role in the regulation of the myogenic response and autoregulation in the cerebral circulation [6, 37, 38]. More recently, emerging evidence indicated that cerebrovascular pericytes expressing alpha-smooth muscle actin (α-SMA) that primarily localize on precapillary arterioles also participate in controlling CBF and maintaining cerebral vascular function with their contractile capability [39-50]. However, whether 20-HETE directly constricts cerebrovascular pericytes and precapillary arterioles and whether it is related to CBF autoregulation in the deep brain cortex have never been studied. The present study seeks to answer these questions using pharmaceutical intervention with a stable 20-HETE analog 20-hydroxyeicosa-5(Z),14(Z)-dienoic acid (WIT003), and a 20-HETE synthesis inhibitor N-Hydroxy-N′-(4-butyl-2-methylphenyl)-formamidine (HET0016) [51-53]. We first identified the expression pattern of CYP4A in cerebral mural cells of the MCAs, penetrating arterioles, and parenchymal arterioles (PAs). We then examined the effects of 20-HETE on the contractile capability of α-SMA^+^cerebral pericytes and primary VSMCs isolated from the MCA of Sprague Dawley (SD) rats. We also studied the effects of 20-HETE on the myogenic response of the PAs and CBF autoregulation in the surface and deep cortex in SD rats.

## 2. Materials and methods

### 2.1. Animals

All experiments were performed using male SD rats purchased from Envigo (Indianapolis, IN). All animals were housed following standard laboratory animal conditions, maintained with free access to food and water ad libitum. The University of Mississippi Medical Center (UMMC) animal care facility is approved by the American Association for the Accreditation of Laboratory Animal Care. The animal protocols were approved by the Institutional Animal Care and Use Committees (IACUC) of the UMMC and followed the NIH’s relevant biosafety policy and guidance.

### 2.2. Cell culture

Primary cerebral VSMCs were isolated from the MCA of 3-week old male SD rats, as described earlier [54-57]. Briefly, the MCAs that were dissected from the brain were placed in ice-cold Tyrode’s solution, containing contained 145 mM NaCl, 6 mM KCl, 1 mM MgCl2.6H_2_O, 50 uM CaCl_2_.2H_2_O, 10 mM HEPES sodium salt, 4.2 mM NaHCO_3_, 10 mM Glucose; pH 7.4. After incubation with papain (22.5 u/mL) and dithiothreitol (2 mg/mL) in the Tyrode’s solution at 37 °C for 15 minutes, the MCAs were centrifuged at 1,500 rpm and resuspended in Tyrode’s solution supplement with elastase (2.4 u/mL), collagenase (250 u/mL), and trypsin inhibitor (10,000 u/mL). The digested vessels were centrifuged, and the pellet of VSMCs was resuspended in the Dulbecco’s Modified Eagle’s Medium (DMEM, Thermo Scientific, Waltham, MA) containing 20% fetal bovine serum and 1% penicillin/streptomycin. The cells were seeded into a 6-well plate pre-coated with Cell-Tak Cell and Tissue Adhesive (354242, 3.5 µg/cm^2^; Corning Inc. Corning, NY) for stocking cells for later experiments or freeze in liquid nitrogen. Early passages (P_2_ - P_4_) of primary VSMCs were used in all experiments.

Human brain microvascular pericytes (HBMVPs) were purchased from Angio-proteomie (cAP-0030, Boston, MA). Early passages (P_3_-P_4_) of HBMVPs were seeded in a cell culture plate pre-coated with Cell-Tak Cell and Tissue Adhesive and incubated in pericyte growth medium (cAP-09, Angio-proteomie) for the following experiments. The purity of pericytes has been validated by staining with several widely accepted markers in our recent study, and > 95% of these cells express α-SMA [50].

### 2.3. Validation of CYP4A expression in cerebral VSMCs and HBMVPs

The expression of CYP4A in cerebral VSMCs and HBMVPs was determined using immunocytochemistry as we described previously [50, 54-56, 58]. Briefly, primary cerebral VSMCs and HBMVPs were seeded on 4-well Nunc™ Lab-Tek™ II Chamber Slide (154526, Thermo Scientific) per-coated with Cell-Tak Cell and Tissue Adhesive. The cells were fixed with 3.7% paraformaldehyde (PFA) and subsequently blocked with 1% bovine serum albumin (BSA) after permeabilization with 0.1% Triton-100 (Sigma-Aldrich, St. Louis, MO). The cells were then co-incubated with anti-α-SMA (1: 300; A2547, Sigma-Aldrich) and anti-CYP4A (1: 200; 299230, Daiichi Pure Chemicals, Tokyo, Japan) primary antibodies in a blocking solution at 4 °C overnight, followed by incubation with Alexa Fluor 555 (1: 1,000; A-31570, Thermo Fisher Scientific, Waltham, MA) and Alexa Fluor 488 (1: 1,000; A-11055, Thermo Fisher Scientific)-labeled secondary antibodies at room temperature. The slides were then coverslipped with antifade mounting medium containing 4′,6-diamidino-2-phenylindole (DAPI; H-1200, Vector Laboratories, Burlingame, CA). Images were captured using a Nikon C2_^+^_ laser scanning confocal head mounted on an Eclipse Ti2 inverted microscope (Nikon, Melville, NY) with a 20 X objective and 2 X digital zoom (total magnification of 880 X). Experiments using primary VSMCs and HBMVPs were repeated 3-4 times in triplicates.

### 2.4. Cell Contraction Assay

Cell contractile capability was compared in primary VSMCs and HBMVPs treated with WIT003 or HET0016 following the protocol provided by a collagen gel-based cell contraction assay kit (CBA-201, Cell Biolabs, San Diego, CA). Briefly, VSMCs or HBMVPs were harvested and resuspended in DMEM or pericyte growth medium at a density of 2 × 10^6^ cells/mL. The cells were then mixed with collagen gel working solution on the ice at a ratio of one in four in volume. The cell-gel mixture solution (0.5 mL, containing 2 × 10^5^ cells) was placed in each well of a 24-well plate and incubated for 1 hour at 37 °C for collagen polymerization. The cells were supplied with an appropriate medium (1 mL) and incubated for 24 hours to develop the contractile force at 37 °C in a 5% CO_2_ humidified atmosphere. The cells were co-incubated with β-Nicotinamide adenine dinucleotide 2′-phosphate reduced tetrasodium salt hydrate (NADPH; 1 mM; N7505, Sigma-Aldrich) and HET0016 (10 μM) in the presence or absence of WIT003 (10 μM) or 0.2% ethanol in culture medium as a control since the stocking solutions of both drugs were 10 mM in ethanol [51, 52]. After incubation, the stressed matrix was detached from the wall of the plate using a sterile needle to initiate the cell contraction. Changes in the collagen gel area were measured using the equation: contraction (%) = [(Area_well_ – Area_gel_) / Area_well_] X 100% and quantified with Image J software as we previously described [54-57].

HET0016 and WIT003 were synthesized and provided by Dr. John R. Falck (Departments of Biochemistry and Pharmacology, University of Texas Southwestern Medical Center) [53].

### 2.5. Isolation of cerebral arteries and arterioles

Cerebral arteries and arterioles were freshly isolated from 3-month male SD rats following the protocol we described previously [55-57]. Briefly, the rats were euthanized with 4% isoflurane. The MCAs and PAs were dissected in an ice-cold calcium-free physiological salt solution (PSS_0Ca_), containing 1 % BSA, 119 NaCl, 1.17 MgSO_4_, 4.7 KCl, 18 NaHCO_3_, 5 HEPES, 1.18 NaH_2_PO_4_, 10 glucose, 0.03 EDTA (in mM, pH7.4). The Pas in the Virchow-Robin space [59-62] were identified within the MCA territory. A segment of the PA without branches was dissected for the myogenic response. Some vessels were carefully dissected to keep a small piece of the MCA, penetrating arterioles and as many branches as possible on the PAs (down to the capillary levels) for immunostaining to identify CYP4A expression patterns in mural cells on the vessel wall in the MCA territory. A phase-contrast image of the vascular structure of isolated vessels was first obtained using the Lionheart automated live-cell imager (BioTek Instruments, Inc., Winooski, VT) at a magnification of 20 X prior to using the laser scan confocal microscope for high-resolution images.

### 2.6. Identification of CYP4A expression in the cerebral mural cells on the vessel wall in the MCA territory

The freshly isolated vessels were fixed with 3.7% PFA and incubated with mixed primary antibodies of anti-α-SMA (1: 300, A2547, Sigma-Aldrich) and anti-CYP4A (1: 200; 299230, Daiichi Pure Chemicals) in a blocking-permeabilizing-staining solution, containing 10% BSA, 2% Triton X-100, and 0.02% sodium azide, at 4 °C for 24 hours in a free-floating manner as we previously described [56]. After incubation with a mixture of Alexa Fluor 555 (1: 1,000, A-31570, Thermo Scientific) and Alexa Fluor 488 (1: 1,000; A-11055, Thermo Fisher Scientific)-labeled secondary antibodies for 2 hours at room temperature and washed gently with phosphate-buffered saline (PBS), the vessels were transferred with a glass Pasteur pipette and placed on a glass Superfrost Plus Microscope Slide (12-550-15, Thermo Scientific), which allows the vessels electrostatically adhere to the glass without additional coating process. The glass pipettes were pre-rinsed with 1% BSA in PBS to prevent sticking. The vessels were repositioned to display all branches without overlap using a glass capillary (6 in, 1.2 mm OD; 1B120-6, World Precision Instruments, Sarasota, FL) thinned with a micropipette puller (Narishige, Tokyo, Japan) and pre-rinsed with 1% BSA. After applied a DAPI antifade mounting medium with hardening formulation (H-1500-10, Vector Laboratories), the slides were coverslipped, and images were obtained by a Nikon C2^+^ confocal head mounted on an Eclipse Ti2 inverted microscope using a 60 X oil immersion objective (Nikon).

### 2.7. Myogenic Response of the PA

Freshly isolated PAs were cannulated with glass micropipettes (6 in, 1.2 mm OD; 1B120-6, World Precision Instruments) bathed with 37 °C PSS (PSS_0Ca_plus 1.6 mM CaCl_2_ and without EDTA) with 0.2% ethanol (solute for HETE0016 and WIT003) and equilibrated with 21% O_2_, 5% CO_2_, 74% N_2_ in a pressure myography chamber (Living System Instrumentation, Burlington, VT). HET0016 and WIT003 were prepared at stock concentrations of 1 mM and 10 mM in ethanol, respectively [51, 52]. The PAs were pressurized to 10 mmHg for a 30 - 60 min equilibration period to develop a spontaneous tone [63-65]. Inner diameters (IDs) of the PA in response to an increase in transmural pressure from 10 to 30 mmHg with or without HET0016 (1 μM) were recorded using a digital camera (MU 1000, AmScope) with a 20 X objective and an IMT-2 inverted microscope (Olympus, Center Valley, PA) connected with the myography chamber. IDs of the PA in response to HET0016 (1 μM) with or without WIT003 (10 μM) at 30 mmHg were recorded at 5, 10, 15, and 20 min after drug administration. At the end of the experiment, transmural pressure was reset to 10 mmHg. The PAs were washed 6 - 8 times with PSS_0Ca_, and the IDs of the PA were determined at 10 to 30 mmHg as described above.

### 2.8. Autoregulation of CBF

The impact of 20-HETE on autoregulation of surface and deep cortical CBF in SD rats was determined with the treatment of HET0016 following the procedure described previously [16, 55, 56, 66]. Briefly, the rats were anesthetized with ketamine (30 mg/kg) and inactin (50 mg/kg). After the trachea was cannulated, the animals were connected to a ventilator (SAR-830, CWE Inc. Ardmore, PA). CO_2_ level (30-35 mmHg) was monitored and controlled by a CO_2_ Analyzer (CAPSTAR-100, CWE Inc.). The femoral artery and vein were implanted with cannulas for mean arterial pressure (MAP) determination and drug delivery. The rat head was placed in a stereotaxic apparatus (model 900, David Kopf, Tujunga, CA), and a thin translucent cranial window was formed using a low-speed air drill. One fiber-optic probe (91-00124, Perimed Inc., Las Vegas, NV) coupled with a laser-Doppler flowmeter (LDF, PF5010, Perimed Inc.) probe was inserted into the brain up to 1.5 - 2 mm for recording the deep cortical CBF [56, 67]. After sealed the open cranial window with bone-wax, another fiber-optic probe was placed above the area without visible vessels within the cranial windows to determine surface cortical CBF. Physiological baseline MAP and CBF on the surface and deep cortex were recorded, and vehicle or HET0016 (2 mg/kg) was slowly infused via the femoral vein. The drug or vehicle was maintained for 45 mins, and the MAP and CBF were recorded every 15 mins. MAP was adjusted to 100 mmHg as experimental baselines. CBF on the surface and deep cortex were recorded at increased pressures from 100 to 180 mmHg in steps of 20 mmHg by phenylephrine infusion (0.5 - 5 μg/min) via the femoral vein. A steady-state CBF was achieved and recorded by maintaining MAP for 5 mins at each stage. The MAP returned to 100 mmHg by withdrawing phenylephrine to obtain a new baseline LDF. CBF was recorded at each stage of MAP reduced from 100 to 40 mmHg by graded hemorrhage in steps of 20 mmHg.

HET0016 was freshly prepared for every *in vivo* experiment following a protocol that was previously reported [68]. Briefly, a working solution composes of an aliquot of 50 µL of HET0016, dissolved in dimethyl sulfoxide (DMSO) at a concentration of 14 mg/mL, mixed with 950 µL of 30% 2-hydroxy propyl-β-cyclodextrin (HPβCD, prepared in sterile water). Final concentrations of DMSO (5%) and HPβCD (28.5%) in working solution were within maximum tolerated doses as nonclinical vehicles for the rat *in vivo* study that was previously validated in nonclinical safety assessment studies by four research organizations [69].

### 2.9. Statistics

All statistical analyses were performed with GraphPad Prism 8 (GraphPad Software, Inc., La Jolla, CA). Results are presented as mean values ± SEM. Significances between groups in corresponding values in cell contraction assay and CBF autoregulation were compared with a two-way ANOVA for repeated measures followed by a Holm-Sidak *post hoc* test as described previously [66, 70, 71]. The significances in corresponding values of changes in the IDs of the PA and CBF before and after drug treatments were compared using a paired *t*-test. *P* < 0.05 was considered statistically significant.

## 3. Results

### 3.1. CYP4A is expressed in rat cerebral VSMCs and pericytes that express α-SMA

We first confirmed that CYP4A and α-SMA are coexpressed in primary cerebral VSMCs isolated from male SD rats (**Figure 1A**). We also found that CYP4A is expressed in a human cerebral pericyte cell line that is α-SMA positive, consistent with our previous report [50] (**Figure 1B**). No nonspecific fluorescence was detected when applied with Alexa Fluor 555 and 488-conjugated 2^nd^ antibodies only in both VSMCs and pericytes (Data not shown).

**Figure 1.**
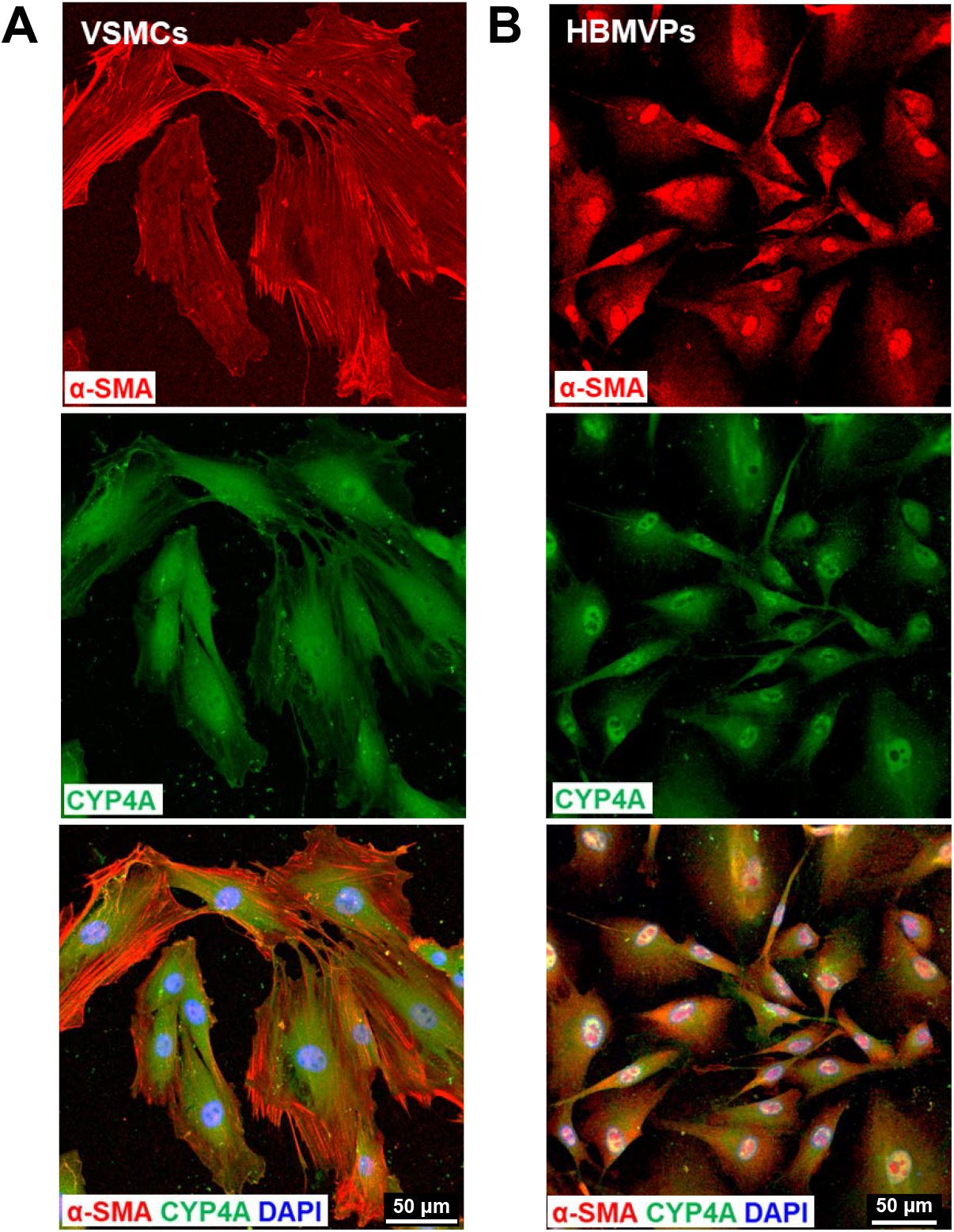
CYP4A is expressed in rat cerebral VSMCs and pericytes that express α-SMA. **A:** Representative images of the expression of CYP4A and α-SMA in primary cerebral VSMCs isolated from the MCA of SD rats. **B:** Representative images of the expression of CYP4A and α-SMA in the HBMVPs. The slides were imaged with a Nikon C2+ confocal head mounted on an Eclipse Ti2 inverted microscope (Nikon) using a 20 X objective and 2 X digital zoom (total magnification of 880 X). VSMCs, vascular smooth muscle cells; α-SMA, alpha-smooth muscle actin; MCA, middle cerebral artery; SD, Sprague Dawley; HBMVPs, human brain microvascular pericytes.

### 3.2. CYP4A expression pattern in the cerebral mural cells on the vessel wall in the MCA territory

A represented phase-contrast image of the vascular structure and organization of isolated vessels in the MCA territory is shown in **Figure 2A**. These arterioles include penetrating arterioles (when branching off from the MCA) and PAs (when entering the brain parenchyma) [57, 60, 62, 72]. The fluid-filled Virchow–Robin space surrounding cerebral microvessels [73, 74] allowed these vessels to be microdissected down to capillary levels without damaging mural cells on the vessel wall.

**Figure 2.**
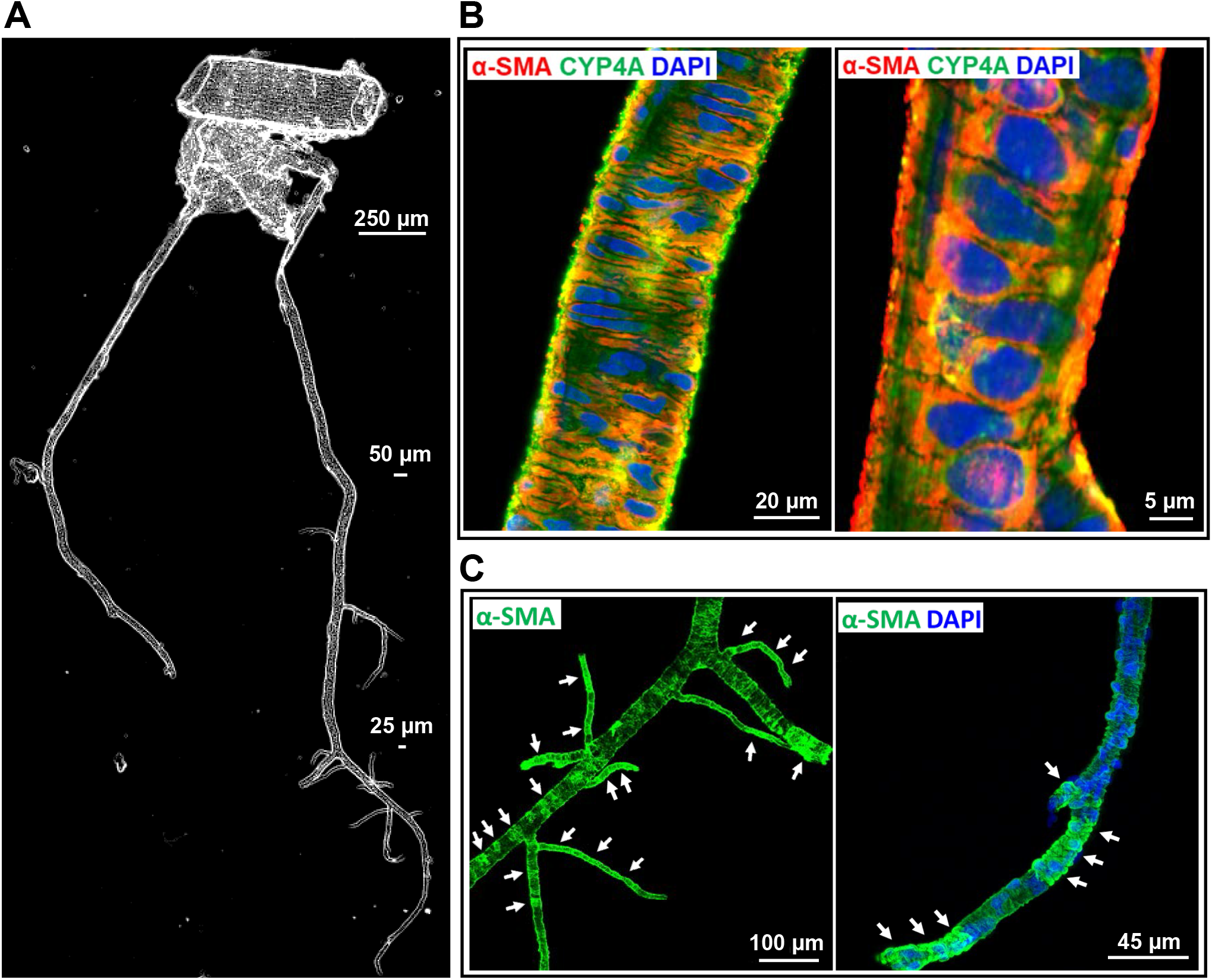
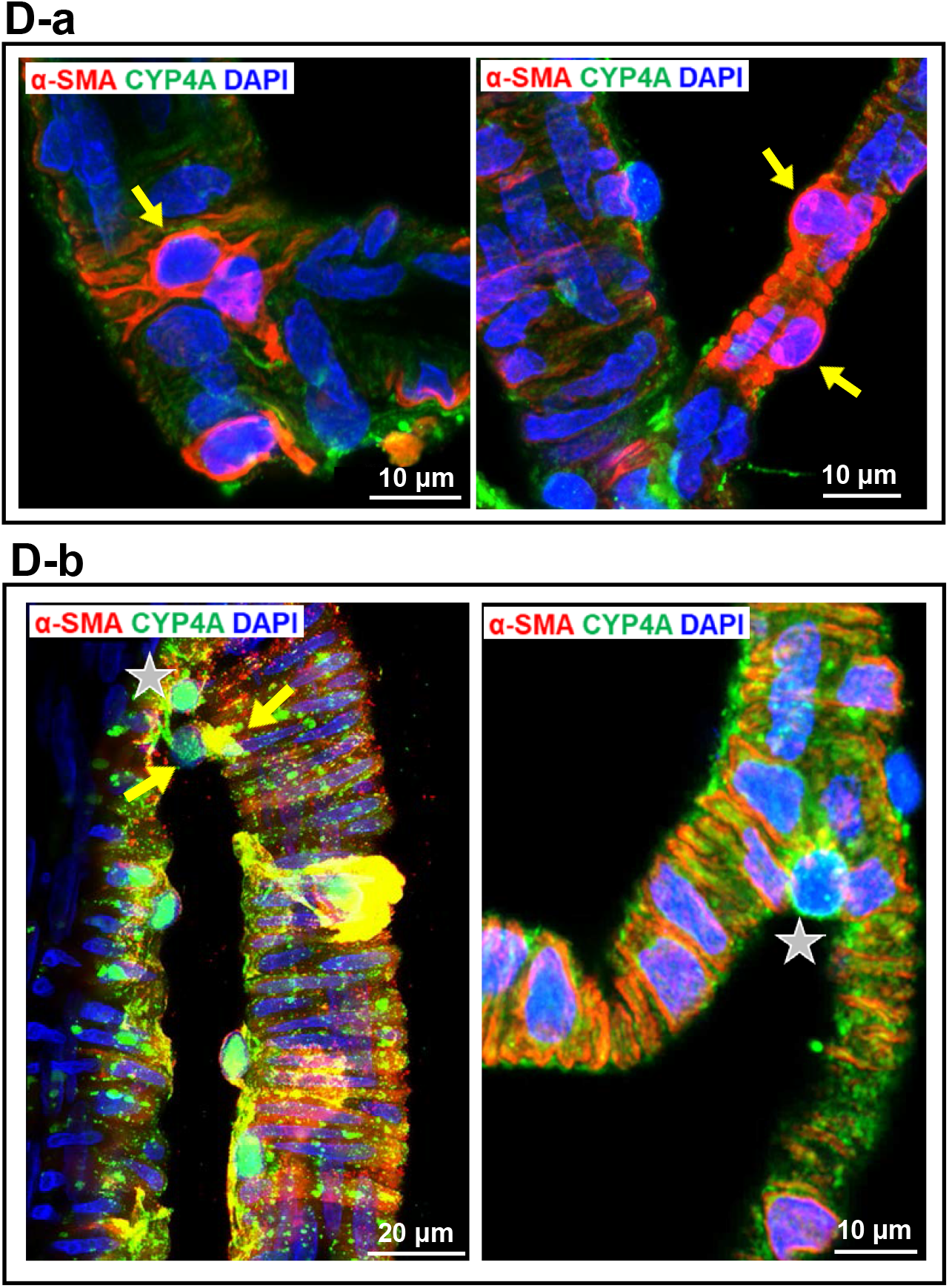
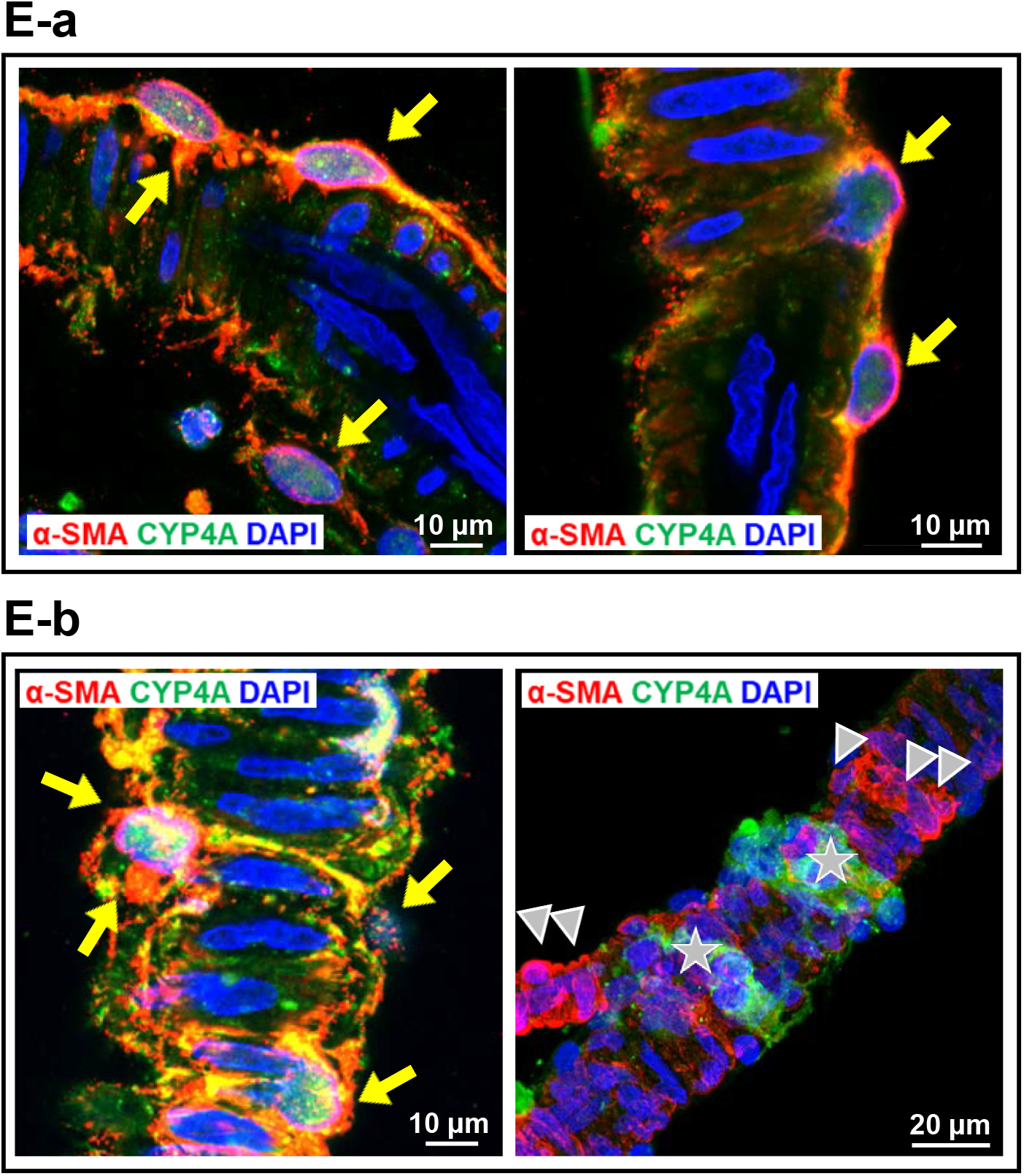
CYP4A expression pattern in cerebral VSMCs and pericytes on the vessel wall in the MCA territory in isolated rat cerebral vasculature. **A:** Representative phase-contrast image of the vascular structure and organization of isolated vessels from SD rats in the MCA territory. **B:** Representative images of the expression of CYP4A and α-SMA in VSMCs in the arterioles. **C:** The expression of α-SMA was detected in mural cells in the arterioles and capillaries in isolated rat cerebral vasculature without a clear defined terminus. **D:** Representative images of the expression of CYP4A and α-SMA in ensheathing or mesh pericytes on the sidewall (a) and junctions of the arterioles and capillaries (b). **E:** Representative images of the expression of CYP4A and α-SMA in thin-strand pericytes longitudinal along with the sidewall (a) and wrapping around the arterioles and capillaries (b). Arrows (white) indicate α-SMA expressed vascular areas (C). Arrows (yellow) indicate CYP4A and α-SMA coexpressed pericytes (D, E). Arrow-heads indicate α-SMA positive but CYP4A negative expressed pericytes. Pentagrams indicate α-SMA negative but CYP4A positive expressed pericyte (D-b) or cell clusters (E-b). Vessel sizes are indicated with size bars. VSMCs, vascular smooth muscle cells; MCA, middle cerebral artery; SD, Sprague Dawley; α-SMA, alpha-smooth muscle actin.

We found that CYP4A and α-SMA were coexpressed in VSMCs of the arterioles downstream of the MCA of male SD rats (**Figure 2B**). CYP4A was also found expressed in all three subtypes of cerebral pericytes, previously defined as ensheathing, mesh, and thin-strand pericytes in mice [39-41]. The expression of α-SMA was found in the arteries, arterioles, and capillaries (**Figure 2C**). Strong coexpression of α-SMA and CYP4A was detected in most ensheathing and mesh pericytes on the wall of arterioles (ID ∼ 20 µm) and capillaries (ID < 10 µm) (**Figure 2D-a**). However, not all mesh pericytes at arteriolar and capillary junctions expressed α-SMA (**Figure 2D-b**). Two types of thin-strand pericytes were found on the wall of arterioles and capillaries: long thin processes longitudinally coursed along the vessels (**Figure 2E-a**) or wrapped encompass the vessels (sphincters) (**Figure 2E-b**); most of these pericytes both expressed α-SMA and CYP4A at the arterioles, but heterogeneity was found at the capillary levels (**Figure 2E-b**). Three to four vessel clusters from four rats were studied. No nonspecific fluorescence was detected when the vessels were incubated Alexa Fluor 555 and 488-labeled 2^nd^antibodies only; however, weak autofluorescence was detected but only at the artery and arteriole levels (Data not shown).

### 3.3. Impact of 20-HETE on cerebral VSMC and pericyte contractile capability

The effects of HET0016 and WIT003 on cerebral VSMC contraction are presented in **Figure 3A**. Gel size was reduced to a lesser extent in HET0016-treated compared with NADPH-treated (control) cells (17.6 ± 0.4 % vs. 28.3 ± 0.5 %), indicating loss of contractile capability after blockade of 20-HETE. In contrast, WIT003 restored VSMC constriction in HET0016 treated cells with a reduced gel size by 29.7 ± 1.0 %. Similarly, as presented in **Figure 3B**, the HBMVPs treated with HET0016 without or with WIT003 exhibited a weaker or stronger contractile capability, respectively, compared to control cells incubated with NADPH only, as the gel sizes decreased by 21.5 ± 1.0 % and increased by 31.6 ± 0.6 %, respectively, compared to 28.6 ± 0.9 %.

**Figure 3.**
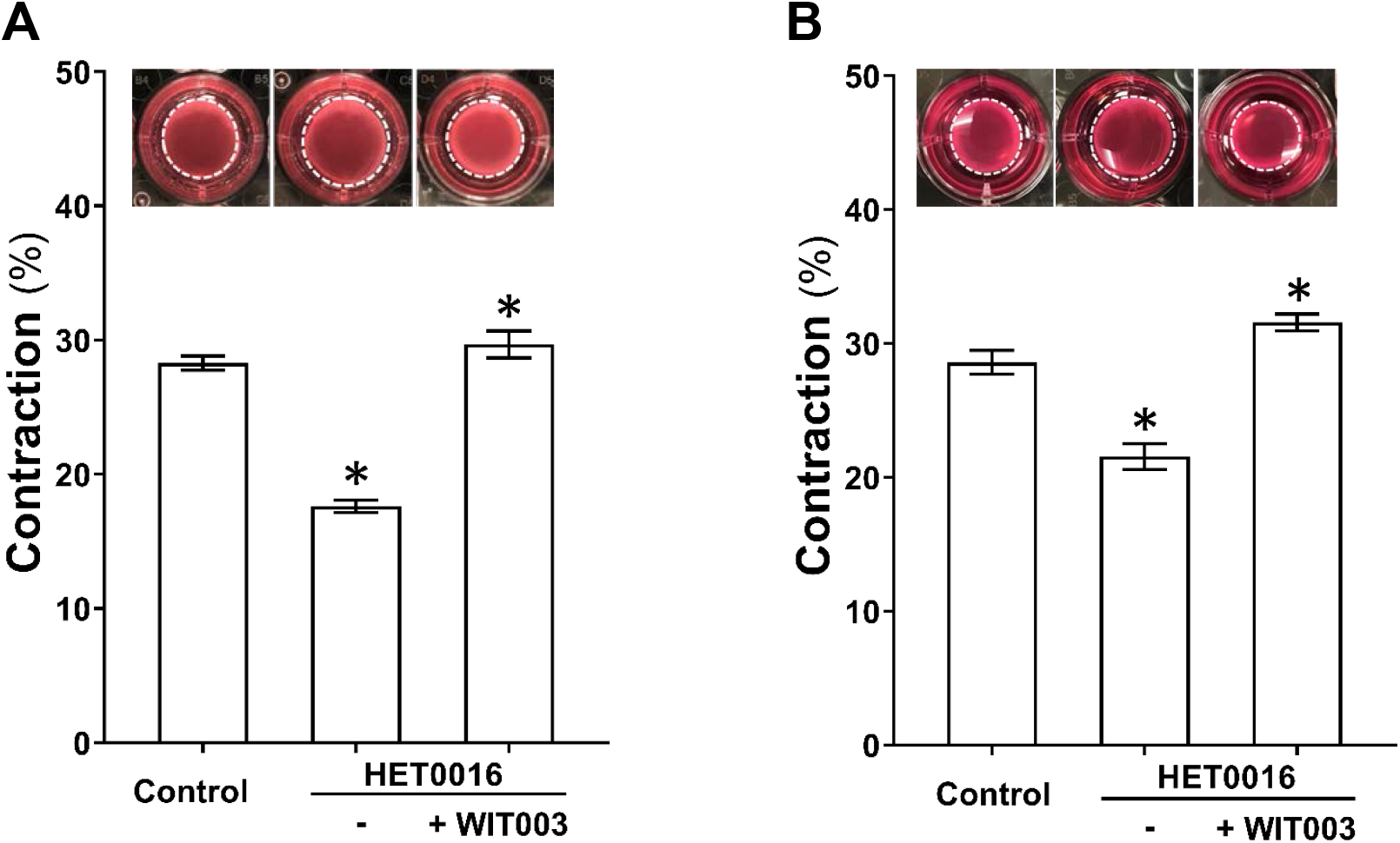
Impact of 20-HETE cerebral VSMCs and pericytes contractile capability. **A:** Comparison of the contractile capability of primary VSMCs isolated from the MCA of SD rat treated with HET0016 in the absence or presence of WIT003. **B:** Comparison of the contractile capability of HBMVPs treated with HET0016 in the absence or presence of WIT003. The inserts are representative images and the white dotted circles represent the gel area after stimulation. Experiments were repeated 3-4 times in triplicates. * indicates *P* < 0.05 from the corresponding values in drugs-treated cells versus controls. VSMCs, vascular smooth muscle cells; MCA, middle cerebral artery; SD, Sprague Dawley; HET0016, N-Hydroxy-N′-(4-butyl-2-methylphenyl)-formamidine; WIT003, 20-hydroxyeicosa-5(Z),14(Z)-dienoic acid; HBMVPs, human brain microvascular pericytes.

### 3.4. Impact of 20-HETE on the myogenic response of the PA

Myogenic response of the PA in SD rats in response to HET0016 and WIT003 were compared. As presented in **Figure 4A**, the PA constricted by 10.2 ± 0.9 % and 17.8 ± 1.5 % when perfusion pressure was increased from 10 to 20 and 30 mmHg, respectively, indicating an intact myogenic response. Administration of HET0016, a 20-HETE synthesis inhibitor, abolished the pressure-induced vasoconstriction of the PA and the vessels dilated by 13.7 ± 4.9 % and 24.1 ± 1.2 %, respectively when perfusion pressure was elevated over the same range. In another group, the PA exhibited maximal dilation by 34.0 ± 3.3 % at 20 min time point after treatment with HET0016 at 30 mmHg. The vasodilation evoked by HET0016 was rescued by WIT003, a 20-HETE analog, and the PA constricted maximally by 23.3 ± 3.6 % 10 min after at the same perfusion pressure (**Figure 4B)**.

**Figure 4.**
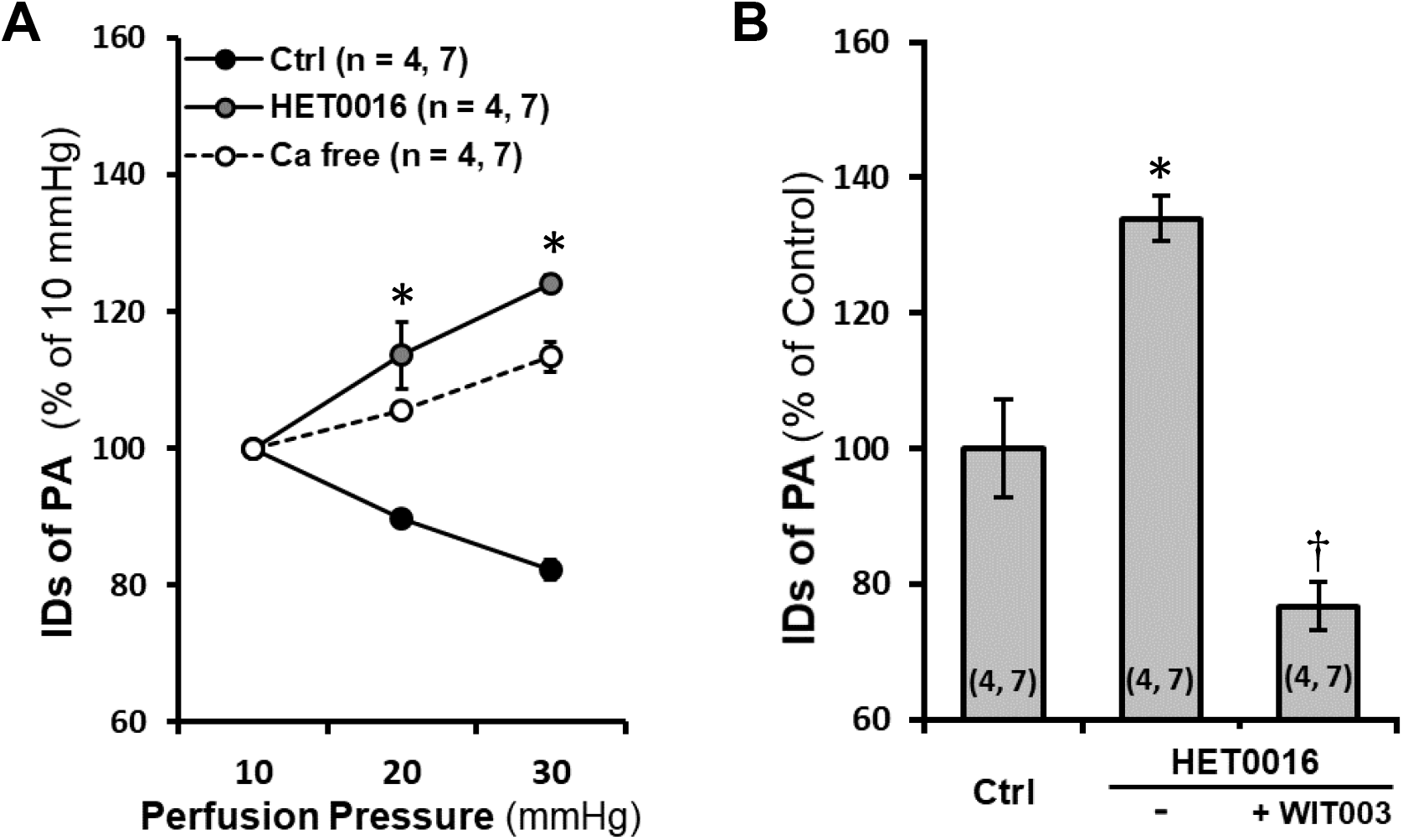
Impact of 20-HETE on the myogenic response of isolated rat Pas. **A:** Comparison of the luminal diameters of the PAs isolated from SD rats in response to increased perfusion pressure with or without HET0016 treatment. Passive diameters were recorded in calcium-free solution. **B:** Comparison of changes in luminal diameters of the PAs isolated from SD rats in response to HET0016 in the absence or presence of WIT003. Mean values ± SEM are presented. Numbers in parentheses indicate the number of animals and vessels studied. * indicates *P* < 0.05 from the corresponding values in control group. † indicates *P* < 0.05 from the corresponding values of HET0016-treated PAs in the absence or presence of WIT003. PAs, parenchymal arterioles; SD, Sprague Dawley; HET0016, N-Hydroxy-N′-(4-butyl-2-methylphenyl)-formamidine; WIT003, 20-hydroxyeicosa-5(Z),14(Z)-dienoic acid.

### 3.5. Impact of 20-HETE on autoregulation of CBF in the superficial and deep cortex

Surface and deep cortical CBF under physiological baseline MAP and CBF autoregulation in response to HET0016 were compared. MAP was not changed in rats when intravenously infused with vehicle or HET0016 for 45 min (data not shown).

Surface cortical CBF did not change in the vehicle-treated group, but it significantly increased by 21.5 ± 5.2 % at 30 min and 25.3 ± 5.5 % at 45 min in rats after HET0016 administration (**Figure 5A)**. Deep cortical CBF also increased significantly by 14.6 ± 3.5 % and 16.9 ± 2.6 %, respectively, in HET0016-treated rats at 30 and 45 min after drug administration (**Figure 5B**).

**Figure 5.**
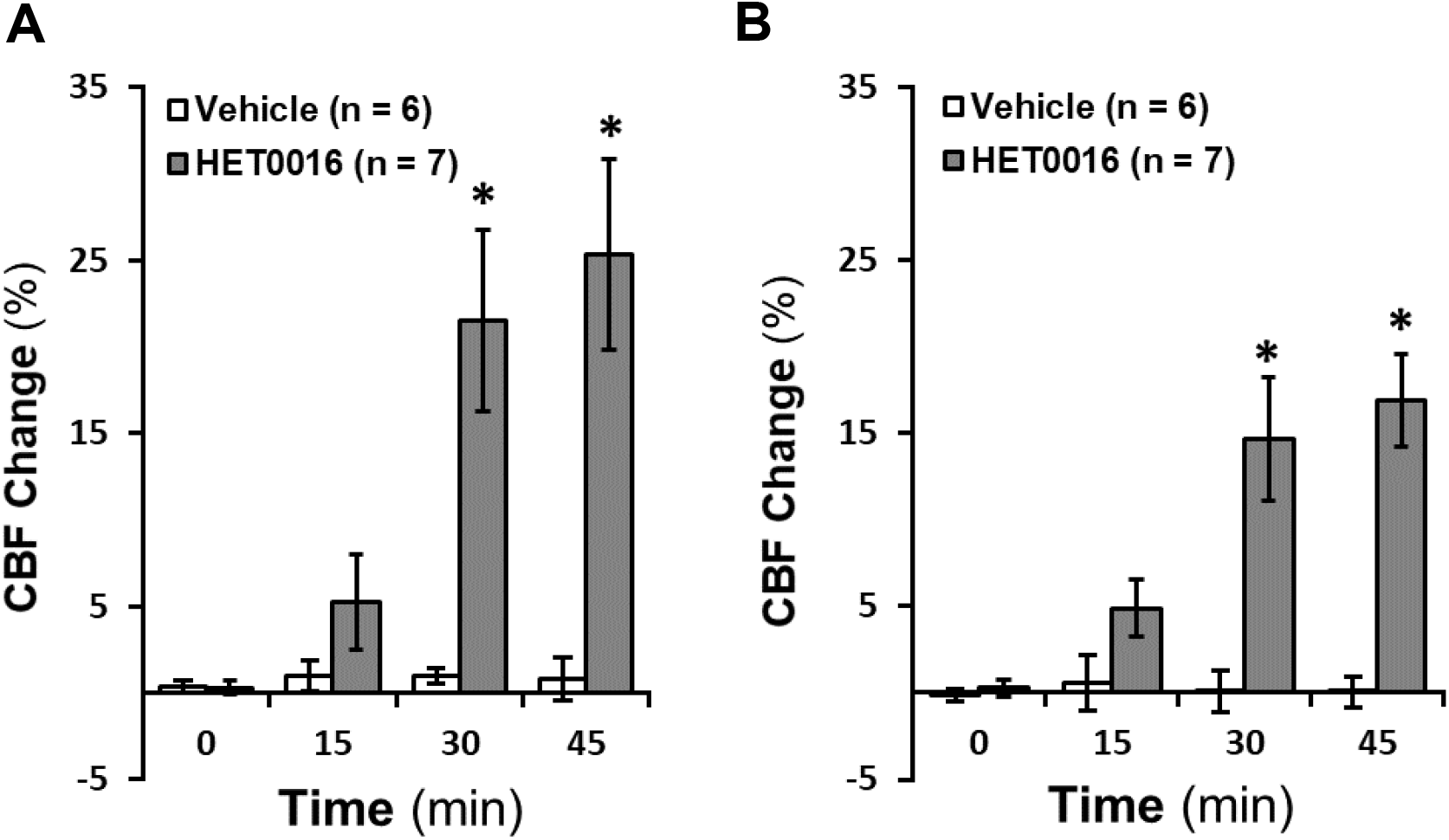
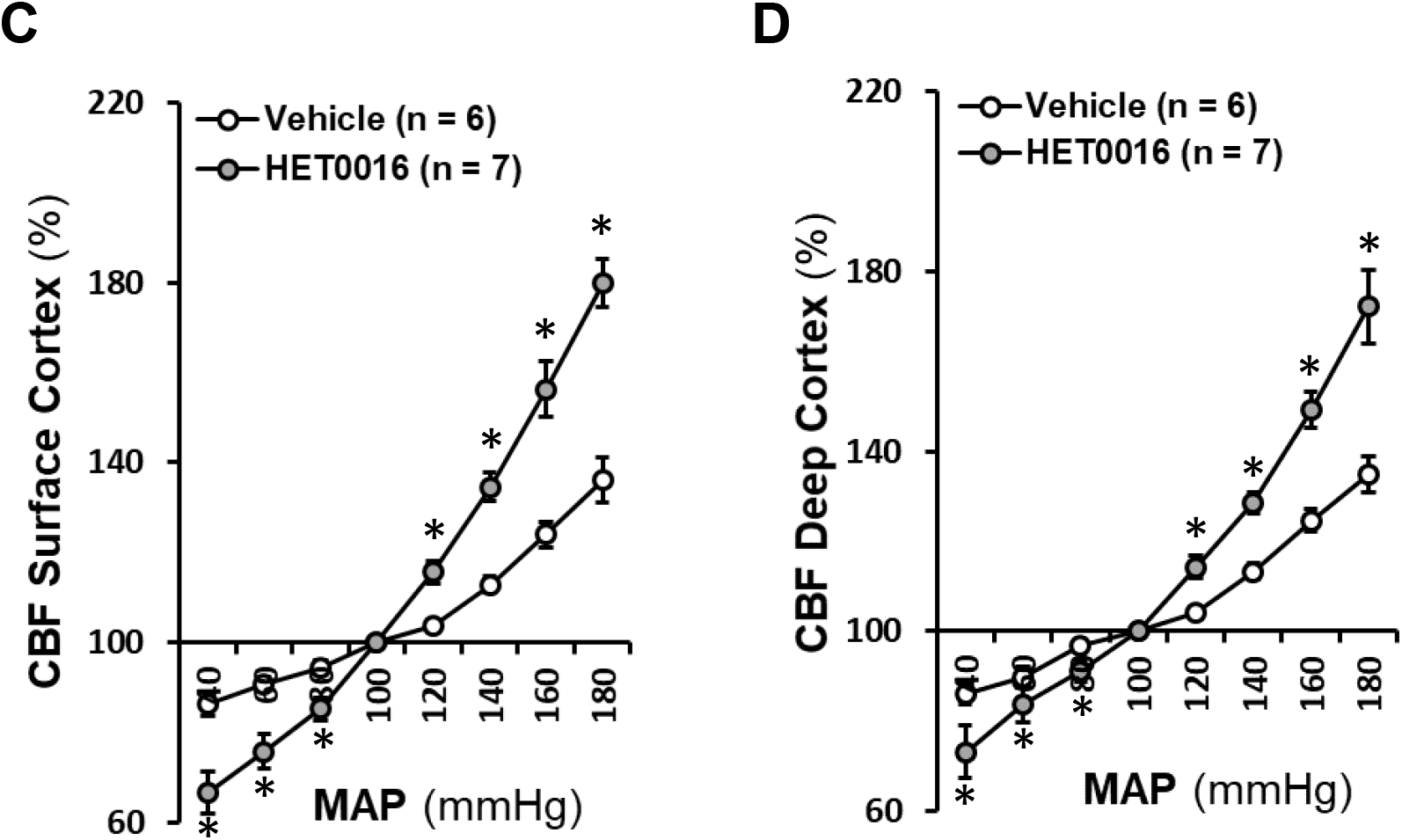
Impact of 20-HETE on CBF autoregulation in rat superficial and deep cortex. **A:** Comparison of rat surface cortical CBF changes in response to HET0016 with time. **B:** Comparison of rat deep cortical CBF changes in response to HET0016 with time. **C:** Comparison of CBF autoregulation in rat superficial cortex. **D:** Comparison of CBF autoregulation in rat deep cortex. Mean values ± SEM are presented. Numbers in parentheses indicate the number of animals studied. * indicates *P* < 0.05 from the corresponding values in HET0016-treated versus -untreated rats. CBF, cerebral blood flow; HET0016, N-Hydroxy-N′-(4-butyl-2-methylphenyl)-formamidine.

As depicted in **Figure 5C** and **5D**, surface and deep cortical CBF autoregulation in 3-month-old male SD rats infused with vehicle were similar as we recently reported [55, 56] with CBF increasing by 12.9 ± 1.7 % and 13.3 ± 1.9 %, respectively, when MAP was elevated from 100 to 140 mmHg. Surface and deep cortical CBF in control rats increased by 36.2 ± 5.0 % and 34.9 ± 3.9 %, respectively, when pressure was increased beyond the autoregulatory range at 180 mmHg; and decreased by 13.6 ± 2.8 % and 13.8 ± 2.6 %, respectively, at 40 mmHg. In contrast, surface and deep cortical CBF in HET0016-treated rats was increased by 34.6 ± 3.2 % and 28.5 ± 2.3 %, respectively, at 140 mmHg; increased by 79.8 ± 5.4 % and 72.1 ± 8.2 %, respectively, at 180 mmHg; and decreased by 33.4 ± 4.6 % and 26.8 ± 5.7 %, respectively, at 40 mmHg.

## Discussion

Emerging evidence over 30 years in human and animal studies demonstrates that 20-HETE exhibits diverse effects in renal and vascular function and plays a pivotal role in hypertension and cardiovascular disease [1, 4-6, 75, 76]. 20-HETE is a product of arachidonic acid catalyzed by the enzymes of CYP4As and CYP4Fs in VSMCs, ECs, pericytes, astrocytes, neurons, and podocytes [31-36]. In the kidney, it inhibits sodium reabsorption and activity of Na^+^/K^+^-ATPase in the proximal tubule (PT) and thick ascending limb of Henle (TALH) [3, 5, 77], and activates transient receptor potential canonical 6 in podocytes contributing to the modulation of glomerular function [36]. In the vasculature, 20-HETE production is inhibited by superoxide, nitric oxide, and carbon monoxide via activation of potassium channels [78]. On the other hand, 20-HETE envokes vasoconstriction and enhances vascular tone by inhibiting potassium channels [13, 79], activating transient receptor potential [80, 81] and calcium channels [82], as well as protein kinase C, Rho- and tyrosine-kinases [83, 84] to enhance intracellular calcium levels.

A substantial body of evidence has suggested that impaired myogenic response and autoregulation in the cerebral circulation is a common pathogenetic neurovascular pathway that plays an essential role in stroke and dementia and in individuals with aging, hypertension, diabetes, and after traumatic brain or spinal cord injuries [56, 85-92]. CBF autoregulation protects the brain from BBB leakage and swelling in response to elevations in blood pressure [60, 93, 94]. One of the predominant regulators of CBF autoregulation is the myogenic response, which evokes vasoconstriction in response to elevations in intraluminal pressure. The myogenic response is an intrinsic property of the VSMCs [95] and pericytes [49]; however, not all cerebrovascular pericytes display constructive capability [46, 96]. Recent studies from the Shih group canonically defined microvascular pericytes in the mouse cortex based on cell morphology, vascular structure and hierarchy, and the α-SMA content [40, 41]. They demonstrated that, unlike VSMCs that have a ring shape with circumferential processes around arteries or arterioles, pericytes have a protruding ovoid cell body. In the mouse cortex, they found an α-SMA terminus at the 1^st^to 4^th^order branches ramifying from the penetrating arterioles. Ensheathing pericytes are localized upstream of the α-SMA terminus, but the mesh and thin-strand pericytes are typically localized downstream of the α-SMA terminus. This observation was recently confirmed [43] that α-SMA-enriched sphincters are localized up to the 4^th^order capillaries in mice cerebral cortex. These sphincter cells exhibited pericyte morphology, which expressed α-SMA. Pericytes that positively express α-SMA regulated CBF, protecting downstream capillaries from elevated perfusion pressure and limiting vasodilation. These cells particularly play a vital role in the regulation of CBF and blood distribution when they localized at the proximal bifurcations of PAs of capillaries [43], which was consistent with a recent report by Gonzales et al., in mice retinal arterioles and capillaries [42, 47] and by our group in cerebral microvessels in rats [31].

Interestingly, results in the present studies in rats are somewhat different from what has been previously reported in mice. We found that there was no clearly defined pattern of the α-SMA terminus in mural cells on the vessel wall in the MCA territory in freshly dissected vessels of 3-month old male SD rats. The expression of α-SMA could be found in mural cells in the arterioles and capillaries. In mice [41], ensheathing pericytes appeared in the PAs and precapillary arterioles upstream of the α-SMA terminus, while mesh pericytes were α-SMA negative that appeared in capillaries only. Notably, ensheathing and mesh pericytes could not be distinguished in rats following such criteria since the “ensheathing” structure at 440 X magnification (**Figure 2C)** became “mesh” when imaging at 2,640 X (**Figure 2D-a)**. Most of these cells expressed α-SMA, spreading from large arterioles to capillaries since there is no clearly defined α-SMA terminus.

However, mesh pericytes that did not express α-SMA were also found, mostly at the arteriolar and capillary junctions (**Figure 2D-b**). Another interesting finding in rats is that the thin-strand pericytes, defined in mice studies, were also expressed α-SMA and localized on the wall of arterioles and capillaries. These pericytes exhibited long thin processes longitudinally coursed along the vessels as in mice. We also found these pericytes wrapped encompass the vessels, similar to the sphincters but predominantly localized in capillaries (**Figure 2E**), consistent with our previous report [31]. However, heterogeneity was found at the capillary levels in terms of the coexpression of α-SMA and CYP4A, although they were detected in most of these pericytes at the arterioles (**Figure 2E-b**).

We found that CYP4A was expressed in VSMCs and α-SMA positive pericytes at arterial levels on the vessel wall in the MCA territory in isolated vessels from rats. The present studies revealed that 20-HETE has a direct impact on cerebral VSMC and pericyte cell contractile capability. Inhibition of 20-HETE production reduced cell constriction in both cell types. These effects were rescued by WIT003. We also found that inhibition of 20-HETE production with HET0016 abolished pressure-induced constriction in the PA of SD rats. The vasodilation effects by HET0016 was rescued by WIT003 in these vessels. These results are in line with our early studies using Dahl S rats, a salt-sensitive hypertensive rat model with a genetic deficiency in the production of 20-HETE and the expression of CYP4A [15]. Dahl S rats exhibited reduced myogenic response of the MCA and poor CBF autoregulation on the surface of the brain cortex [16]. The impairments of cerebral vascular hemodynamics were rescued by knocking in wildtype *Cyp4a1* onto the Dahl S transgenic background. CYP4A expression and 20-HETE production were globally enhanced, including cerebral vasculature in this model [16]. The current *in vivo* studies demonstrated that inhibition of 20-HETE synthesis diminished CBF autoregulation at both surface and deep cortex, further confirming the direct impacts of 20-HETE on cerebral VSMC and pericyte cell contractile capability play an important role in the regulation of CBF.

Poor CBF autoregulation has been found to accelerate detrimental cerebrovascular consequences in aging and various disease [56, 85-89, 97-100]. We have reported that this impairment could transmit elevated pressure to penetrating arterioles and contribute to cerebral vascular disease in another aging hypertensive rat model [101]. We also have reported that the pathogenetic mechanism in CBF autoregulation was found in the aging diabetic model [56], rats with genetic defects or genetically modified in genes regulating vascular function [57, 66, 102]. Compared with the VSMCs on the MCA, pericytes on the PA exhibit an increased calcium channel activity, the lack of large-conductance Ca^2+^-activated potassium (BK) channel, and greater myogenic tone at physiological pressures [56, 72]. PAs protect downstream capillaries from BBB leakage and swelling by enhancing resistant quality in microvessels in the deep cortex [103]. Contractile pericytes at capillary junctions in mice retinal vasculature regulate branch-specific blood flow by receiving hyperpolarizing signals propagating through the capillary network induced by potassium [42].

Nevertheless, heterogeneity of cerebrovascular pericytes in morphology, localization, α-SMA content could be due to differences in species and approach. However, it is most likely due to the diverse effects of cerebrovascular pericytes and their plasticity in the brain under different genetic and environmental conditions [104, 105]. Pericytes regulate CBF, redistribute capillary blood flow, maintain the BBB integrity, and play many other roles such as angiogenesis and phagocytosis [45, 96, 106-109]. Pericyte constriction in response to a thromboxane A2 analog displayed a nonsimultaneous “on and off” operation [44]. The heterogeneity of the coexpression of α-SMA and CYP4A in pericytes at arteriolar and capillary levels suggests CYP4A in α-SMA negative pericytes at the neurovascular unit may play a different role, which may involve pericyte-endothelia crosstalk to maintain BBB integrity. Indeed, it is under-explored how to define brain pericytes into subtypes in different species under different physiological and pathological conditions, how pericyte heterogeneity and plasticity play different roles, and how pericytes produce, receive and propagate signals to crosstalk with other brain cell types.

## Conclusions

Results from the current study indicated, for the first time, that 20-HETE-promoted CBF autoregulation is associated with enhanced α-SMA positive cerebrovascular pericyte contractility in rats. This observation is associated with the coexpression of CYP4A and α-SMA in cerebral VSMCs and most pericytes. Our findings provide a novel insight into possible mechanisms by which inactivating variants of 20-HETE producing enzymes have been found in hypertension, stroke, and AD patients [10, 18-29].

## Funding

This study was supported by grants AG050049, AG057842, P20GM104357, and HL138685 from the National Institutes of Health.

## Declaration of competing interest

None.

## Notes

### Competing Interest Statement

The authors have declared no competing interest.

